# Discovery of a Natural Microsporidian Pathogen with a Broad Tissue Tropism in *Caenorhabditis elegans*

**DOI:** 10.1101/047720

**Authors:** Robert J. Luallen, Aaron W. Reinke, Linda Tong, Michael R. Botts, Marie-Anne Félix, Emily R. Troemel

## Abstract

Microbial pathogens often establish infection within particular niches of their host for replication. Determining how infection occurs preferentially in specific host tissues is a key aspect of understanding host-microbe interactions. Here, we describe the discovery of a natural microsporidian parasite of the nematode *Caenorhabditis elegans* that has a unique tissue tropism compared to other parasites of *C. elegans*. We characterize the life cycle of this new species, *Nematocida displodere,* including pathogen entry, intracellular replication, and exit. *N. displodere* can invade multiple host tissues, including the epidermis, muscle, neurons, and intestine of *C. elegans*. Despite robust invasion of the intestine very little replication occurs there, with the majority of replication occurring in the muscle and epidermis. This feature distinguishes *N. displodere* from two closely related microsporidian pathogens, *N. parisii* and *N*. sp. 1, which exclusively invade and replicate in the intestine. Comparison of the *N. displodere* genome with *N. parisii* and *N*. sp. 1 reveals that *N. displodere* is the earliest diverging species of the *Nematocida* genus and devotes over 10% of its genome to a single species-specific gene family that may be mediating host interactions upon infection. Altogether, this system provides a convenient whole-animal model to investigate factors responsible for pathogen growth in different tissue niches.

## Introduction

Pathogens infect host organisms and then often establish themselves within a particular niche of the host environment in order to replicate [1-3]. This niche usually resides within a particular cell type or tissue, and is commonly referred to as cellular or tissue tropism. The mechanisms responsible for tissue tropism are broad and potentially multifactorial, and can involve features such as access, specific receptor/ligand interactions, pathogen competence for growth in particular tissue niches, and/or host defense [4-8]. Understanding the mechanistic and evolutionary basis for tissue tropism is key to understanding host/pathogen interactions and discovering therapeutics to prevent pathogens from causing disease.

Microsporidia represent a large phylum of obligate intracellular pathogens related to fungi, which can infect a diverse array of hosts from protists to humans [9-12]. They have features consistent with having adapted to proliferate exclusively within the host cellular environment, including greatly reduced genome size and the loss of true mitochondria [13]. Different species of microsporidia display a range of different tissue tropisms. For example, the microsporidian species *Encephalitozoon cuniculi*shows a broad tissue tropism in humans and is able to infect the liver, brain, kidneys, skin, and gastrointestinal tract, while *Enterocytozoon bieneusi* mainly infects the enterocytes of the small intestine [14]. Studying tissue tropism in higher animals can be confounded by the complexity of the host body plan, making it difficult to comprehensively describe the tissues that are subject to infection in vivo. In some cases, tropism is implied from in vitro studies based on cell types that are infected and may not reflect the true tropism within the live animal [15, 16].

The nematode *Caenorhabditis elegans* is a tractable, whole-animal system to study host/pathogen interactions because of its simple body plan and transparency, which facilitates assessment of tissue tropism in vivo. Sampling of proliferating populations of *Caenorhabditis* nematodes from rotting plant substrates in wild habitats has demonstrated that they are regularly infected by microsporidia [17-19]. Isolates of two closely related species, *Nematocida parisii* and *Nematocida* sp. 1, are thus far the only described microsporidian species in wild *Caenorhabditis* nematodes, and both of these species are fecal/oral pathogens that infect and replicate exclusively in *C. elegans* intestinal cells [9, 18]. The intestinal-trophic nature of *N. parisii* has been well-studied, with all stages of the pathogen being solely observed in the intestine by light, fluorescence, and transmission electron microscopy (TEM) [18]. Additionally, multiple infection-induced changes have been observed in the intestine, including restructuring of the apical cytoskeleton [20] and hijacking of the intestinal recycling endosome pathway for the exit of newly made spores [21].

Here, we report the discovery of a new species of microsporidia found infecting non-intestinal tissues of a wild-caught *C. elegans* animal. Whole genome sequencing and phylogenomic analysis places this new species in the *Nematocida* genus, and we have named it *Nematocida displodere,* based on a bursting mechanism for spore exit. *N. displodere* displays a distinct tropism from the other described *Nematocida* species and has the capacity to initially invade a wide array of tissues and cell types, including the intestine, epidermis, muscle, neurons, and specialized phagocytic cells called coelomocytes, with feeding being required for infection. Interestingly, the majority of intestinal infection fails to replicate. Comparison of the *N. displodere* genome with the other *Nematocida* species shows that despite *N. displodere* having a smaller genome, it contains an enormously expanded species-specific gene family that comprises over 10% of its predicted protein-coding genes, which may explain its distinct infection life cycle. Altogether, we characterize a new species of microsporidia, *Nematocida displodere,* with a broader tissue tropism yet smaller genome than the other *Nematocida* species identified to date, and this provides a convenient model to investigate mechanisms responsible for pathogen growth in different tissue niches.

## Results

### Discovery of a new species of microsporidia that infects a broad range of tissues in *C. elegans*

While sampling for nematodes near the Viosne stream in Santeuil, France, we found a wild-caught *C. elegans* infected by a microbe displaying microsporidian-like features in the head of the animal (S1a Fig). For reference, microsporidian species display certain stereotypical hallmarks in their life cycle [22]. Specifically, infection begins when an extracellular, transmissible spore fires an infection apparatus called a polar tube to deliver a single mononucleated parasite cell called a sporoplasm into the host cell. The sporoplasm then develops into a multinucleate, proliferative stage called a meront, and eventually differentiates into new spores that exit the host cell. The wild-caught *C. elegans* we found had structures that appeared like meronts and spores in an area that is likely the epidermis (S1a Fig). In the lab, we found that this P_0_ adult was able to transmit infection to its progeny, as observed by the appearance of large, meront-like structures and spores in recipient animals (Fig 1a-b). These later stages of infection were seen along the anterior/posterior axis of the animal, including the head (Fig 1a-b), the mid-body (S1c, Fig 1c), and the tail (Fig 1c). The majority of animals showed these later stages of infection in the body wall of *C. elegans,* which is the outer tube of the animal distal to the pseudocoelomic space that includes the epidermis, muscle, and neurons [23]. By contrast, tissues in the intestine (Fig 1a) and gonad remained for the most part symptom-free as observed by light microscopy. Consistent with a lack of infection in the gonad, we did not find that infection was vertically transmitted because the eggs from a bleached population of heavily infected animals developed into a population that remained uninfected for multiple generations (n=100 animals analyzed by Nomarski and n=100 animals analyzed by fluorescent in situ hybridization, FISH, over 4 months at 15°C).

**Fig 1.**
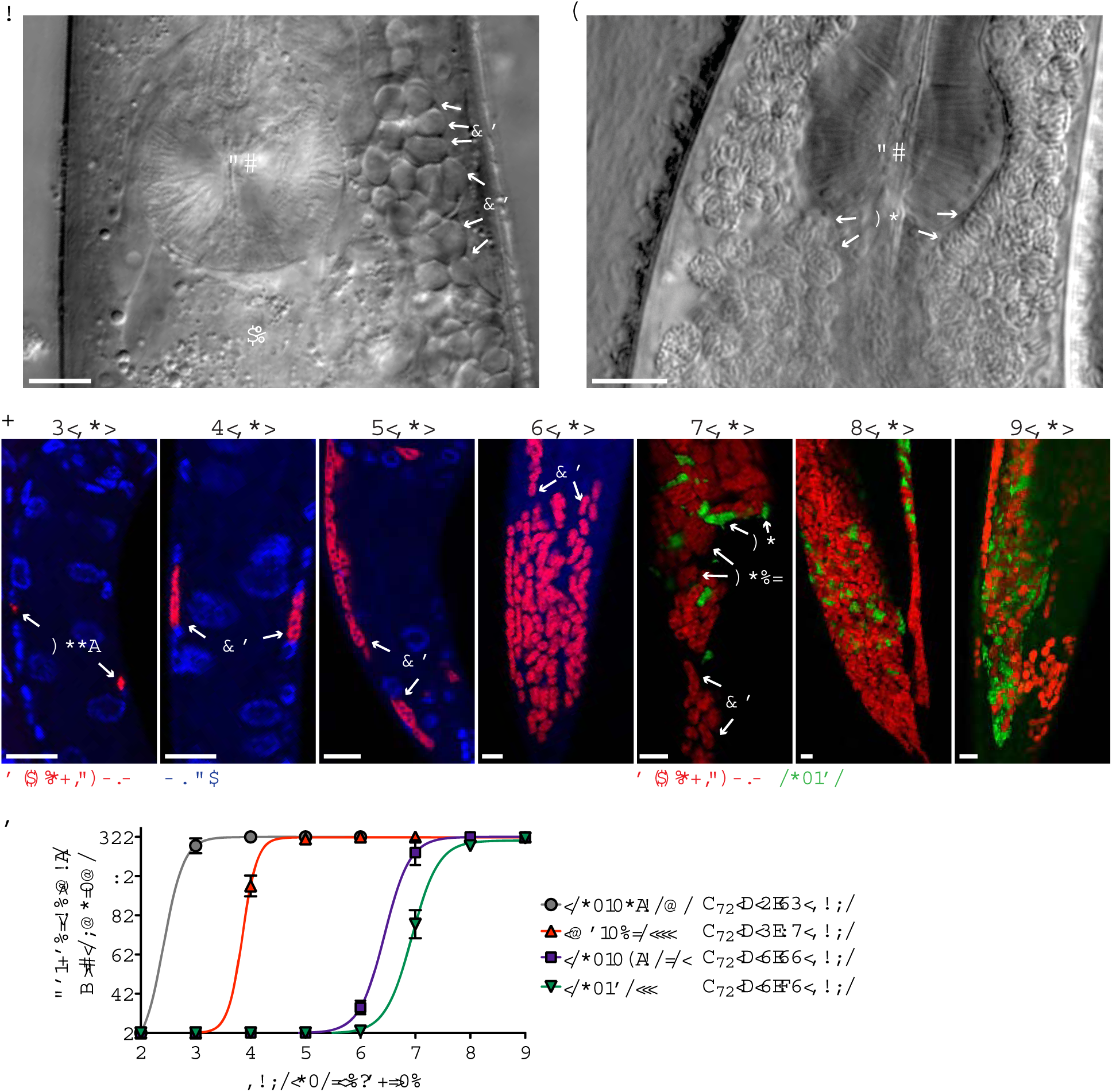
A new microsporidian species that infects *C. elegans*. (a) Infected head region of a live *C. elegans* animal from strain JU2807 (derived from the wild-isolated P_0_ animal, see Supplementary Fig 1) showing a large group of structures that appear to be meronts (*Me*) adjacent to the pharynx (*Ph*) and intestine (*In*). (b) Infected head region with the area adjacent to the pharynx filled with spores (*Sp*). (c) The mid-posterior to tail region of N2 *C. elegans* infected with *N. displodere* visualized by FISH to the parasite rRNA (red), DAPI for nuclei (blue), and direct yellow 96 (DY96, green) staining of parasite spores from 1 dpi to 7 dpi at 15°C. Animals were at the L2 larval stage at 1 dpi, L3 stage at 2 dpi, L4 stage at 3 dpi, and adult stage at 4-7 dpi. Sporoplasms (*Sppl*), meronts (*Me*), sporonts (*Spnt*), and spores (*Sp*) are indicated. Scale bars are 10 *μ*m. (d) Quantification of symptoms of *N. displodere* infection over time at 15°C with N2 animals infected as starved L1 larvae at T_0_. Sporoplasms are mononucleated structures, meronts are multinucleated structures, and sporoblasts are rounded, mononucleated structures stained by FISH (see c above). Spores are oblong DY96-stained structures in infected animals. Fifty animals were quantified for each replicate at each time point, and data points indicate the mean and SD from four replicates across two experiments. Each symptom was fit to a Boltzmann sigmoidal curve (R square > 0.99 for each curve), and the time to 50% of the animals exhibiting symptoms (T_50_) is shown.

We confirmed this pathogen as a new species of microsporidia in the *Nematocida* genus based on whole genome sequencing and phylogenomic comparison (see below), and we named this new species *Nematocida displodere*. To characterize the infection life cycle of *N. displodere* we labeled the pathogen with a FISH probe targeting the small ribosomal subunit RNA (rRNA). We synchronized wild-type N2 *C. elegans* animals, infected them with *N. displodere* spores and then observed them for the main hallmarks of microsporidia infection with this FISH probe. These hallmarks include mononucleated sporoplasms observed at 1 day postinfection (dpi), multinucleate meronts from 2-4 dpi, and sporoblasts (pre-spores) and spores at 5 dpi, which eventually fill up a large proportion of the animal by 7 dpi (Fig 1c). We quantified the percent of animals exhibiting the symptoms of each stage of infection using FISH from 1 to 7 dpi (Fig 1d), and found that 100% of animals in a population exhibited replicative forms of infection by 2 dpi, and 100% exhibited spores by 7 dpi. Thus, similar to the intestinal-trophic *N. parisii, N. displodere* appears to efficiently infect an entire *C. elegans* population on a plate in a laboratory setting [18].

To more closely examine the *N. displodere* life cycle in *C. elegans,* we conducted transmission electron microscopy (TEM) analysis on infected animals. When compared to uninfected animals (Fig 2a), we observed numerous structures by TEM that looked like distinct stages of microsporidia infection. These structures include large, multinucleate cells that are likely the proliferative meront stage (Fig 2b) and groups of mononucleate cells that likely correspond to the sporont stage of microsporidia, which are thought to be able to undergo further divisions (Fig 2c) [24]. Additionally, groups of cells were seen with nascent microsporidian spore structures that likely represent sporoblasts, which do not undergo any further divisions before becoming spores (Fig 2d-e). Finally, darker, more fully differentiated spores were seen (Fig 2f-g). From the sporont to the spore stage of *N. displodere* we observed cross-sections of polar tube coils (the specialized infection apparatus of microsporidia). In spores that appeared fully developed, a maximum of five polar coils were observed per cell (seen in 19 of 70 TEM cross-sections of spores), with two coils on one side and three coils on the other (Fig 2c, g). Thus, *N. displodere* appears to undergo stereotypical developmental features of microsporidia, as assessed by TEM.

**Fig 2.**
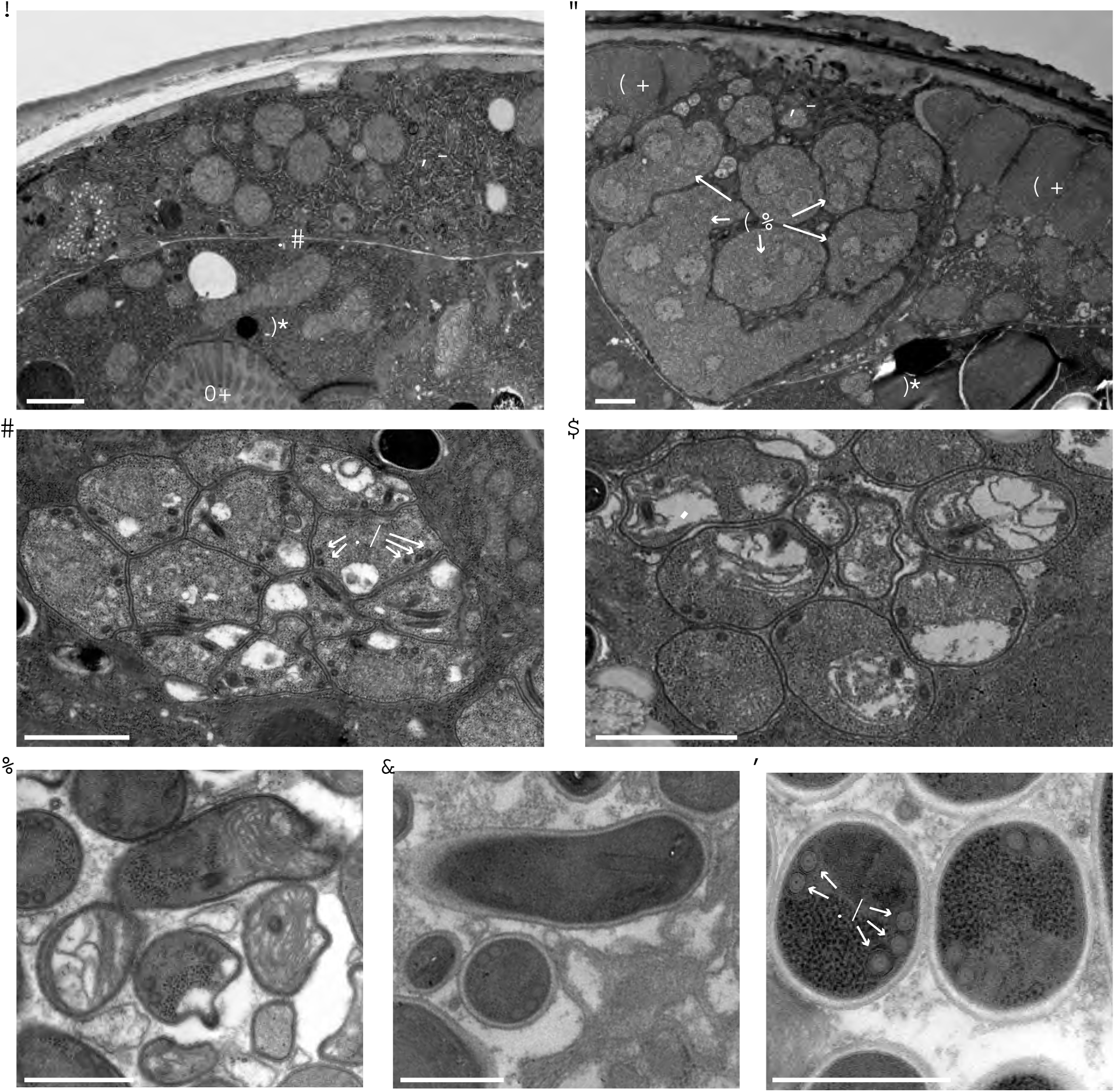
Transmission electron micrographs of *N. displodere*-infected C. elegans. (a) Cross-section of an uninfected adult with the epidermis (*Ep*) and intestine (*In*) shown, separated by the pseudocoelom (*Pc*). The intestinal lumen (*Lu*) is indicated, (b) Cross-section of an *N. displodere-'mtecteti* adult at 6 dpi with large meronts (*Me*) presumably in the epidermis, adjacent to two flanks of the body wall muscle (*Mu*), (c) Large associated cluster of *N. displodere* sporonts with nascent polar tube coils (*PT*) in an infected animal at 8 dpi. (d) Groups of nascent spores, presumably sporoblasts, in an 8 dpi animal, (e) Groups of nascent spores, presumably sporoblasts, in an 8 dpi animal. (f) Longitudinal view of a spore in an 8 dpi animal. (g) Crosssectional view of spores in an 8 dpi animal, with five polar tube coils. Scale bars are 1 μm (a-d) and 500 nm (e-g).

### *N. displodere* can invade multiple *C. elegans* tissues, but preferentially prcliferates and differentiates in the epidermis and muscle

Our observations of *N. displodere* infection by light and electron microscopy indicated that meronts and spores were predominantly in non-intestinal tissues, suggesting a different tropism than *N. parisii,* which exclusively infects the intestine. To more carefully compare the tissue tropism of these two species, we co-infected N2 animals with *N. displodere* and *N. parisii* and found that indeed these two closely related microsporidian species infect distinct areas of the animal (Fig 3a). Next, to determine the range of tissues in which *N. displodere* can proliferate, we infected a panel of *C. elegans* strains that express GFP in distinct tissue types, and then looked for multinucleate meronts at 3 dpi by rRNA FISH, and newly differentiated spores at 5 dpi with a chitin-binding dye that stains spores (Direct Yellow 96 – DY96). Using this approach, we found the epidermis, body-wall muscle, and neurons had *N. displodere* meronts (Fig 3b) and newly-formed spores (Fig 3c). Additionally, we occasionally saw meronts of *N. displodere* in epidermal seam cells and coelomocytes (S2 Fig), but we did not observe new spores in these cells at later time points. In some cases, *N. displodere-infected* cells appear to become larger than corresponding uninfected cells, as can be seen in the neurons (Fig 3b, *bottom*) and seam cells (S2a Fig). Additionally, we found that multiple tissues can be infected in the same animal. For example, meronts were found both inside and outside of the muscle in one animal (Fig 3c, *middle*).

**Fig 3.**
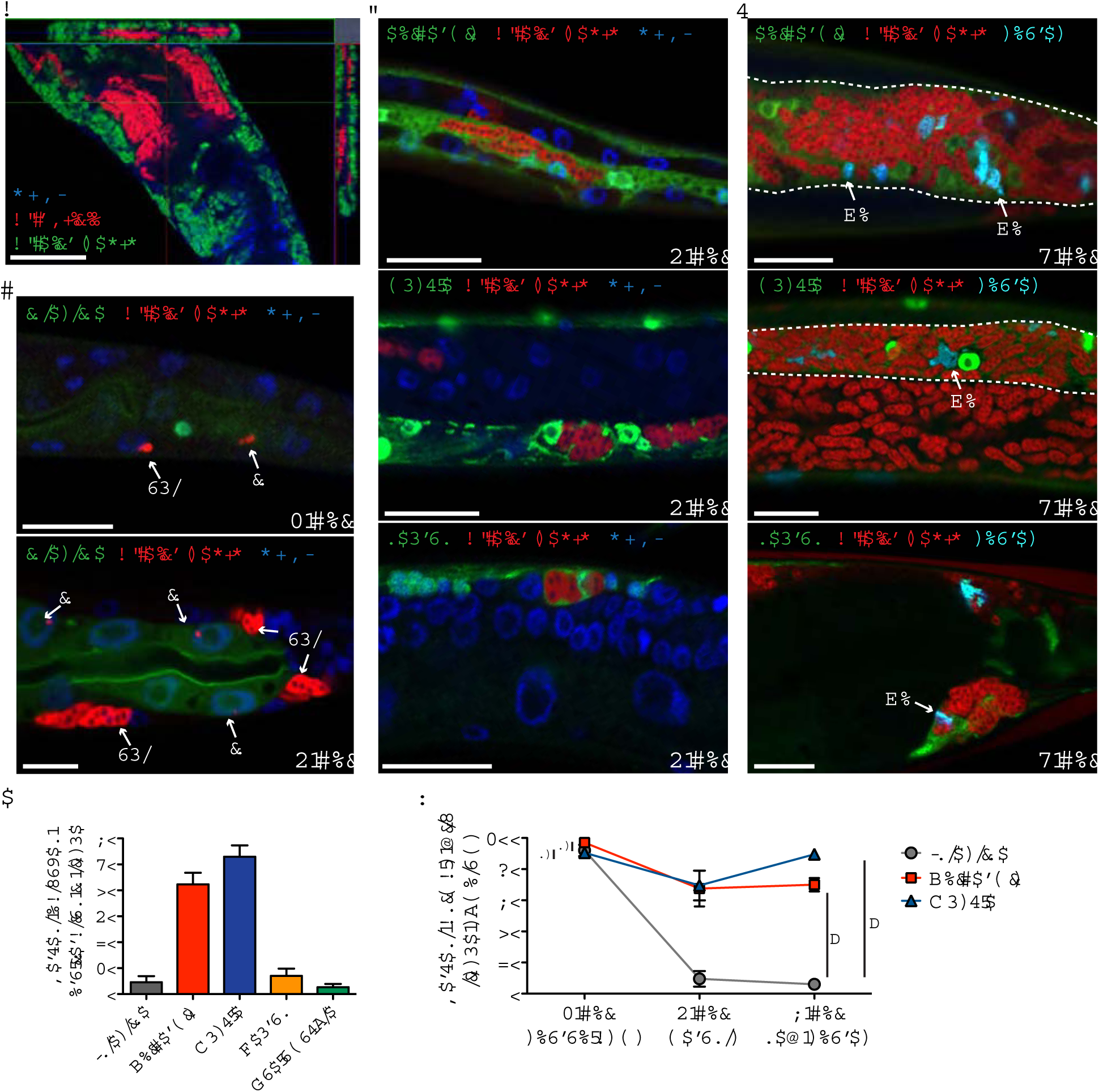
*N. displodere* infects multiple tissues but shows preferential proliferation in non-intestinal tissues. (a) The anterior region of a *C. elegans* animal co-infected with *N. displodere* (green) and *N. parisii* (red), visualized by FISH using species-specific rRNA probes and DAPI (blue). This image was captured by confocal microscopy with a single z-plane represented in the main inset, and orthogonal views of the x‐ and y-planes on the top and right insets, respectively, which show a cross-sectional view of the captured z-stacks within those planes. Scale bar is 50 *μ*m. (b) *C. elegans* tissue-specific GFP-expression strains in the epidermis (*top*), body wall muscle (*middle*), and neurons (*bottom*), were infected with *N. displodere* and imaged at 3 dpi by FISH and DAPI. The neuron infected was in the ventral nerve cord (*bottom*). (c) Tissue-specific GFP strains were infected and imaged at 5 dpi with FISH and DY96 to stain spores (*Sp*). GFP-positive tissues that are difficult to see due to heavy infection are outlined with dashed white lines. The neuron infected was in the pre-anal ganglia (*bottom*). Scale bars are 20 *μ*m. (d) The mid-body of the *C. elegans* intestinal GFP-expression strain infected with *N. displodere* at 1 dpi (*top)* and 3 dpi (*bottom*). Infection events are labeled as either inside (*in)* or outside (*out)* of the GFP-labeled intestine. Scale bar is 10 *μ*m. (e) The tissue distribution of proliferating *N. displodere* infection was analyzed at 3 dpi, and was calculated individually in each *C. elegans* tissue-expression strain as the percent of FISH-stained meront clusters occurring in the GF-positive tissues compared to the total number of events throughout the animals. Data are represented as the mean with SD of four replicates across two experiments, with a total of 50 animals counted for each replicate. (f) A comparison of the percent of animals infected in the specified GFP-positive tissue at three time points at which the three main stages of *N. displodere* infection occur, with sporoplasms analyzed at 1 dpi, meronts at 3 dpi, and new spores at 6 dpi. Each time point was calculated individually in each *C. elegans* tissue-expression strain as the percent of 50 animals that show a given symptom in the GFP-positive tissues. Data are represented as the mean with SD of four replicates across two experiments (ns=not significant, comparing intestine to muscle (p=0.55) or intestine to epidermis (p=0.11) at 1 dpi; *p=0.03 comparing intestine to muscle and comparing intestine to epidermis at 6 dpi, twotailed Mann-Whitney test).

We observed far fewer animals infected in the intestine compared to the muscle and epidermis at 3 and 5 dpi, although a small fraction of animals did show meronts and newly made spores in the intestine (S3 Fig). Interestingly, the intestine was frequently invaded with *N. displodere* sporoplasms at 1 dpi, but was rarely infected with multinucleate meronts at 3 dpi (Fig 3d). This observation suggests that *N. displodere* can invade this tissue, but often fails to proliferate there. We quantified this observation by counting the total number of meront clusters in an animal at 3 dpi and then calculating the percentage of those clusters that are in a particular GFP-labeled tissue. This analysis was performed separately for each tissue in its respective tissue-specific GFP expression strain, with each cluster of meronts assumed to have formed from a single invasion event. In this manner, we found that only about 5% of multinucleate meronts were found in the intestine, whereas 42% were in the epidermis, 53% in the muscle,7% in the neurons, and 3% in the coelomocytes (Fig 3e). Although the infections in each tissue were quantified separately using individually marked *C. elegans* strains, this approach appears to provide a good estimate of the overall tissue distribution of meronts at 3 dpi, because the total percent of pathogen in these five tissues adds up to about 100%.

We next looked at other time points to quantify the tissues in which *N. displodere* could invade to deliver sporoplasms, proliferate into multinucleate meronts, and differentiate into spores We infected the intestinal, muscle, and epidermal GFP expression strains for 1, 3, or 6 days and counted the fraction of animals displaying a given *N. displodere* stage at that time point in the GFP-labeled tissue. While greater than 90% of animals were initially infected with sporoplasms in either the intestine, epidermis, or muscle at 1 dpi, very few of these invasion events appeared to proliferate and differentiate in the intestine, with less than 10% of animals exhibiting meronts in the intestine at 3 dpi and 6% exhibiting new spores in the intestine at 6 dpi (Fig 3f) By contrast, greater than two-thirds of animals showed meronts and new spores in either the muscle or the epidermis at these later time points. Together, these results suggest that *N. displodere* can initially invade multiple tissues in *C. elegans,* including the intestine, but shows preferential proliferation and differentiation in the epidermis and muscle.

### *C. elegans* infection with *N. displodere* is dependent on feeding

We next investigated how *N. displodere* is able to access the host environment to invade host tissues. Invasion via the epidermis or the intestine after feeding are the only two routes described for pathogens infecting *C. elegans*[25, 26], and these two tissues represent the largest surface areas with exposure to the environment. To investigate whether *N. displodere* infection of *C. elegans* might occur through feeding or external penetration through the cuticle, we investigated transcriptional responses characteristic of intestinal infection and cuticle damage. First, we analyzed the *C. elegans* intestinal GFP reporter strains for the genes *F29F2.1* and *C17H1.6,* which are highly induced upon infection with both *N. parisii* and another natural *C. elegans* intestinal pathogen, the Orsay virus [27]. Here we found that *N. displodere* infection caused induction of both the *F29F2.1p::GFP* (Fig 4a, *left)* and the *C17H1.6p::GFP* reporter strains (S4 Fig). In fact, *N. displodere* caused a greater degree of induction of these reporter strains than *N. parisii*. By contrast, an epidermal infection/damage reporter strain for *nlp-29,* a gene highly induced upon infection by the fungus *D. coniospora* and epidermal wounding through the cuticle [26, 28], did not show induction after infection by *N. displodere* (Fig 4a, *right). N. parisii* infection also failed to induce *nlp-29,* as has been previously described [18]. The lack of *nlp-29* induction suggests that despite its capacity to infect the epidermis, *N. displodere* is unlikely to cause damage through cuticle disruption, which is known to induce *nlp-29* [26]. These results, together with the fact that *N. displodere* spores were seen in the intestinal lumen soon after infection (S5 Fig), suggest that feeding could be a major route for infection.

**Fig 4.**
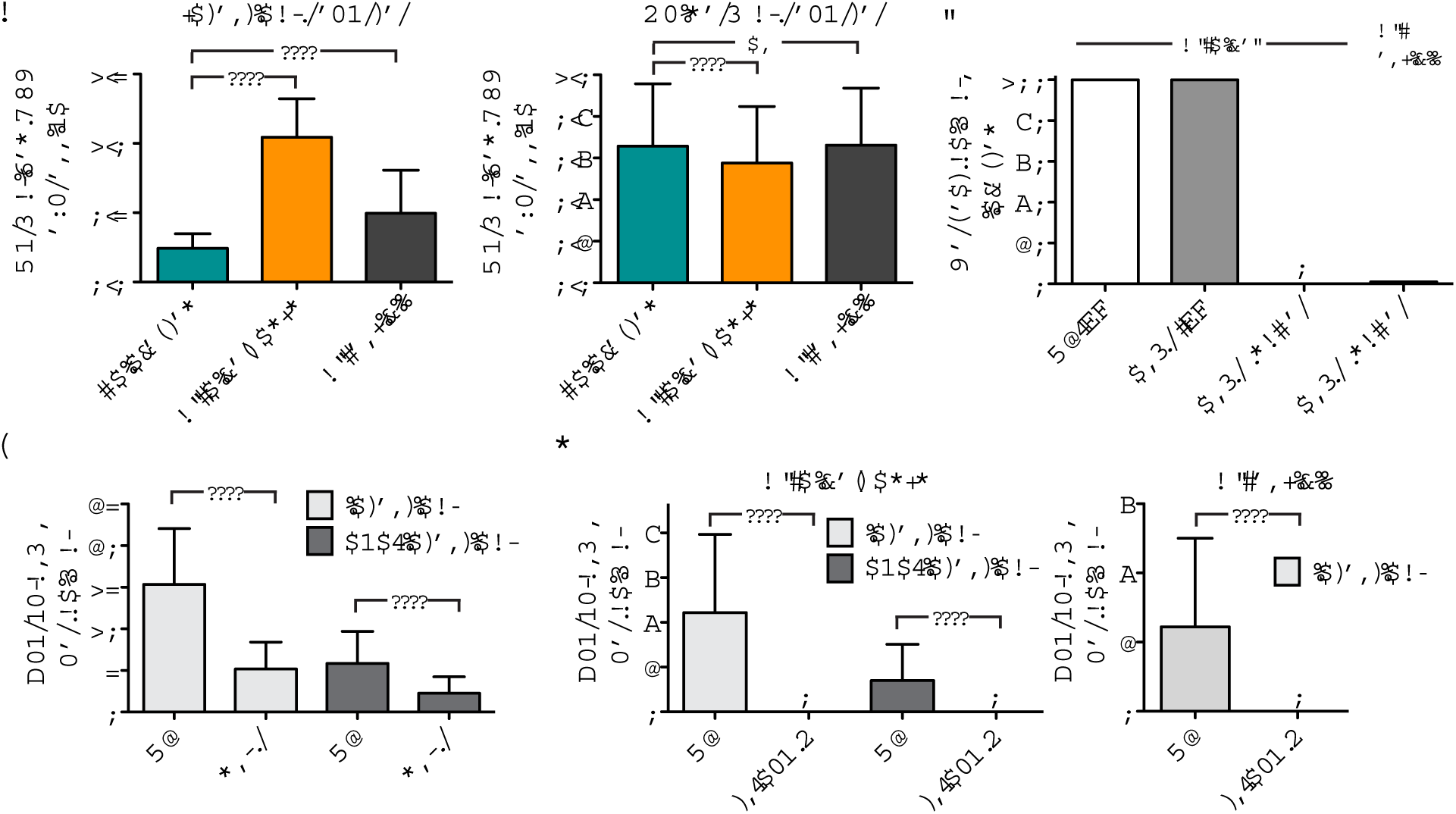
*N. displodere* induces an intestinal response, and host feeding is required for infection. (a) Normalized GFP induction after *N. displodere* or *N. parisii* infection of an intestinal infection reporter strain (ERT54, *left*), and an epidermal infection/cuticle damage reporter (AU189, *right*), as measured by a COPAS Biosort. Experimental replicates were normalized by animal body size for ERT54 or by red fluorescence (*pcol12::dsred*) for AU189. For ERT54, data are represented as mean values with SD from n=882 animals from six replicates across two independent experiments (****p<0.0001, two-tailed Mann-Whitney test). For AU189, due to a batch effect, only data are shown from three replicates in one experiment, with mean values shown with SD from n=900 animals (****p<0.0001, ns=not significant, two-tailed Mann-Whitney test). Data from the other AU189 replicates are shown in the supplement (Supplementary Fig 4). (b) Comparison of *N. displodere* infection of *daf-2(ts)* animals at the L3 stage (maintained at 15°C) or induced to form dauer larvae (maintained at 25°C). As a control, N2 animals were maintained at 15°C and infected with *N. displodere* as L3 animals at 25°C, and *N. parisii* spores were used to infect *daf-2(ts)* dauer larvae. (c) Comparison of the number of invasion events (counted as sporoplasms) occurring in N2 and *eat-2* animals at 1 dpi. Events were counted as either intestinal (co-localizing with intestinal gut) or non-intestinal. Data are represented as mean values with SD from n=75 animals from three independent experiments (****p<0.0001, two-tailed Mann-Whitney test). (d) Comparison of the number of invasion events (counted as sporoplasms) occurring in *dyn-1(ts)* and N2 animals at 30°C for 30 minutes for *N. displodere (left)* and *N. parisii* infection (*right*). Infection events were distinguished as either intestinal or non-intestinal as above. *dyn-1(ts)* animals are paralyzed and cease to feed at the nonpermissive temperature (30°C). Data are represented as mean values with SD from n=80 *dyn-1(ts)* animals and n=50 N2 animals across two independent experiments (****p<0.0001, twotailed Mann-Whitney test).

To further examine if *N. displodere* infects through *C. elegans* feeding, we assessed infection in *C. elegans* strains that have a reduced or eliminated ability to feed. First, we used the temperature sensitive (ts) strain *daf-2(e1368)* [29], which constitutively enters the nonfeeding dauer stage at the restrictive temperature of 25°C. Dauers of *daf-2(ts)* animals grown at 25°C were inoculated with *N. displodere* and showed no infection in any tissues, while 100% of L3 animals of *daf-2(ts)* maintained at a permissive temperature did show infection (Fig 4b). Similar results were seen with *N. parisii* infection. A caveat to these results being evidence for a feeding-based mechanism of infection is that the dauer stage not only ceases feeding, but also develops a tougher cuticle surface.

Next, we used a *C. elegans eat-2* mutant to investigate whether feeding was important for *N. displodere* infection. The *eat-2* mutant shows an approximately 70% reduction in feeding rate compared to wild-type animals, either in the absence or presence of *N. displodere* spores (S6 Fig). Consistent with this feeding defect, we saw a 66% reduction in sporoplasms in the intestine (Fig 4c). We also saw a 61% reduction in sporoplasms in non-intestinal tissues in *eat-2* mutants compared to wild-type animals (Fig 4c). These results, like the dauer results described above, show that feeding is likely a route of entry for both intestinal and non-intestinal tissue infection by *N. displodere*.

Finally, we tested the *C. elegans* temperature-sensitive endocytosis strain, *dyn-1(ts)* which stops feeding and moving at 30°C. We shifted adult N2 and *dyn-1(ts)* animals to 30°C for 2.5 hours and then infected with *N. displodere* for 30 minutes. Even with this short infection time, wild-type animals showed a substantial level of infection, with an average of 4.4 sporoplasms inside of the intestine and 1.4 sporoplasms outside of the intestine (Fig 4d, left). By contrast, the *dyn-1(ts)* mutants showed a complete lack of infection in any tissue of the animals. We saw similar results for *N. parisii* infection (Fig 4d, right), with substantial intestinal infection in wild-type animals, and no infection in *dyn-1(ts)* mutants. As a control, we verified that *dyn-1(ts)* animals were infected by *N. displodere* at the permissive temperature (S7 Fig). Altogether, these data strongly suggest that *C. elegans* feeding is required for the majority, if not all, *N. displodere* infection in susceptible tissues of *C. elegans*.

### *N. displodere* likely accesses non-intestinal tissues from the intestinal lumen through the use of the polar tube

The dependence of non-intestinal infection on feeding raises the question of how these tissues are accessed from the pharyngeal-intestinal lumen. Notably, the muscle, epidermis, and neurons are in the body wall of *C. elegans* and are separated from the intestine by multiple cell membranes and the pseudocoelom [23]. *N. displodere* spores can be seen in the pharyngeal and intestinal lumen soon after infection, but they are never observed in any other locations in the animals at the early time points of 1 hpi and 24 hpi, including in any other tissue, the pseudocoelom, or even inside intestinal cells (see S5 Fig). These observations suggest that the majority of *N. displodere* infection in non-intestinal tissues originates from the lumen of either the pharynx or intestine. Notably, the pharynx of *C. elegans* is coated by its own secreted cuticle [30], and we have never observed invasion events (sporoplasms) anterior to the posterior bulb of the pharynx (0 of 100 infected animals at 1 hpi) (Fig 5a, S8 Fig). In fact, on the occasions in which *N. displodere* sporoplasms are observed anterior to the intestine, they are seen very near the intestinal lumen, which forms a wide lumenal pocket where the pharyngeal valve cells meets the four most anterior intestinal cells (S8 Fig). Together, these observations suggest infection originates not from the pharyngeal lumen, but from the intestinal lumen.

**Fig 5.**
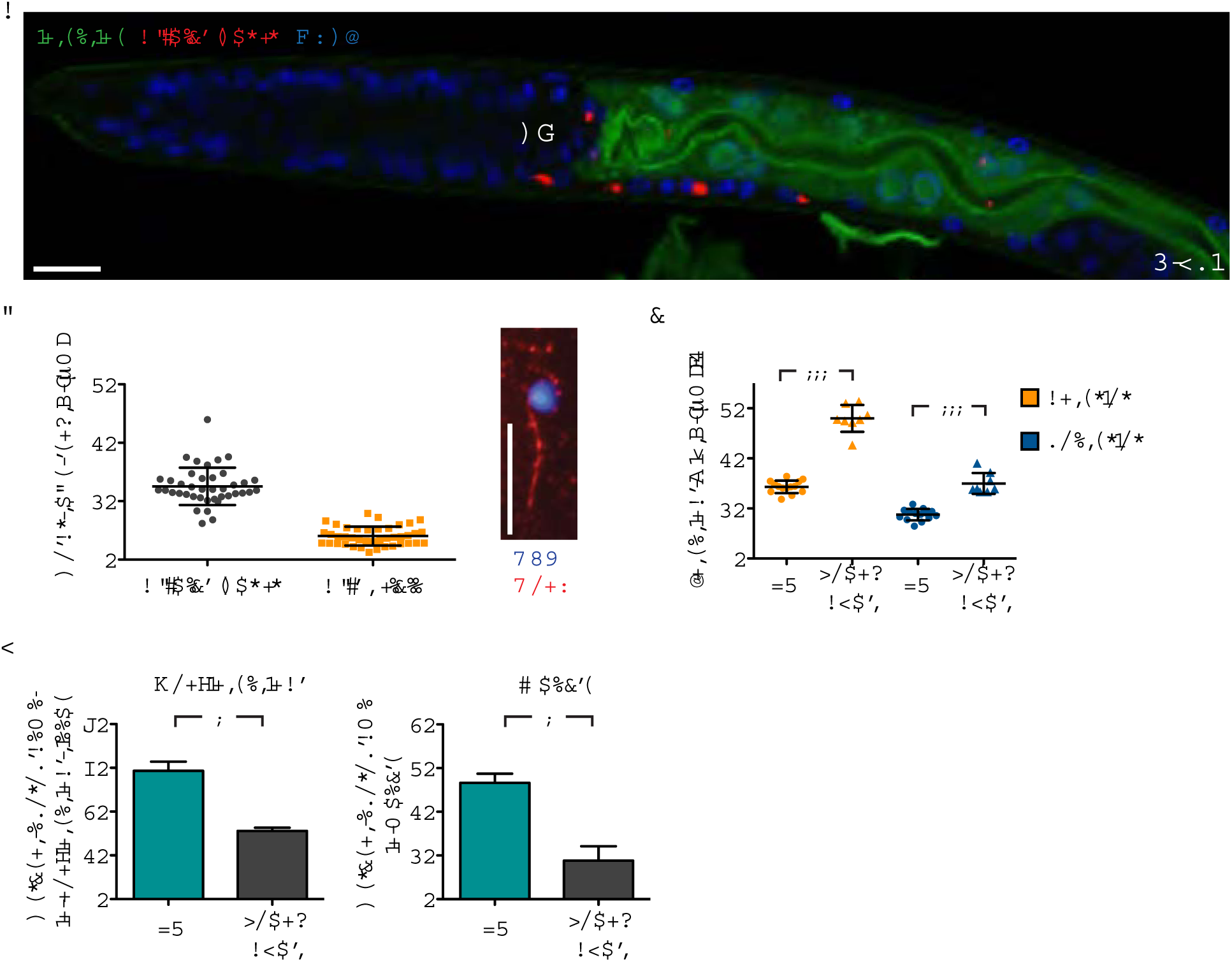
*N. displodere* likely accesses non-intestinal tissues from the intestinal lumen. (a)*C. elegans* intestinal GFP expression strain ERT413 at 1 dpi stained with *N. displodere* rRNA FISH. Sporoplasms are seen inside and outside of the GFP-labeled intestine, in close proximity to the intestine, but never anterior to the posterior bulb (*PB*). Scale bar is 10 μm. (b) Exterior polar tubes associated with a spore were measured for *N. displodere* and *N. parisii* small spores. Each data point represents a measured polar tube, with the line and error bars showing the mean and SD of n=40 for *N. displodere* and n=41 for *N. parisii*. Note polar tubes of *N. displodere* and *N. parisii* were measured with separate techniques on separate occasions. The image (*right)* shows an *N. displodere* spore stained by Calcofluor white (CFW) with the associated polar tube stained by Concanavalin A-rhodamine (ConA). Scale bar is 10 *μ*m.(c) The widths of GFP-labeled intestine from L3 larvae and young adults were measured in the anterior and posterior regions of the animal and halved to estimate the distance from the lumen to the basal lateral side of the intestine. The mean and SD from n=14 L3 animals and n=8 young adults are indicated (***p=0.0002, two-tailed Mann-Whitney test). (d) The tissue distribution of invasion events of *N. displodere* infection (sporoplasms) was analyzed after 30 minutes of infection in L3 larvae versus young adults, and was calculated in the tissue-specific strains expressing GFP in the intestine (*left*)and muscle (*right*). Invasion events were calculated as the percent of FISH-stained sporoplams occurring out of GFP-expressing intestine (*left*) or in GFP-expressing muscle (*right*) compared to the total number of events throughout the animals. Data are represented as mean with SD of four replicates across two experiments, with a total of 25 animals counted for each replicate (*p=0.0286, two-tailed Mann-Whitney test).

Microsporidia invade host cells using a specialized infection apparatus called a polar tube, which is fired upon external stimulus in order to pierce and inject the sporoplasm into the host cell [31]. Although we have been unable to actively observe this invasion process with either *N. displodere* or *N. parisii* in *C. elegans,* we investigated whether it was theoretically possible for the polar tube of *N. displodere* to reach non-intestinal tissues from the intestinal lumen. First, we measured the in vitro length of the *N. displodere* polar tube at 12.55 *μ*m (+/-3.20 *μ*m), and found that these polar tubes were three-fold longer than those of *N. parisii* small spores (measured at 4.03 *μ*m +/-1.61 *μ*m) (Fig 5b). Note that *N. parisii* develops two different sized spores, large and small, with small spores being sufficient for transmitting infection in *C. elegans* [18]. We found that *N. displodere* produced only one observable spore size, measured as 2.38 *μ*m (+/-0.26 *μ*m) long and 1.03 *μ*m (+/-0.18 *μ*m) wide (S9 Fig), which are similar to previous measurements of *N. parisii* small spores (2.18 *μ*m long, 0.8 *μ*m wide) [18].

Next, to estimate if the polar tube of *N. displodere* is long enough to reach non-intestinal tissues from the lumen, we measured the average distance from the lumen to the basolateral side of the intestine in L3 larvae and young adult *C. elegans* using an intestinal GFP expression strain. At the posterior end of the intestine, this distance was measured at 8.8 *μ*m in L3 animals and 15.0 *μ*m in young adults, while at the anterior end it was 14.3 *μ*m in L3 animals and 27.2 *μ*m in young adults (Fig 5c). These measurements are rough estimates, as the lumen of the intestine can have a convoluted path instead of just being a straight line (see Fig 5a, S8 Fig), which results in variability of distances from the lumen to the basolateral side of the intestine. Altogether, however, these data show that the *N. displodere* polar tube is long enough to traverse through an intestinal cell from at least some locations of the lumen, and that these distances are on average smaller in younger compared to older animals.

If infection of non-intestinal tissue is dependent on the polar tube, then we would expect younger animals would have a higher percentage of non-intestinal infection compared to older animals, because the distances from the lumen to the basolateral side of the intestine are shorter in L3 larvae compared to young adults. To test this model, we compared the percent of intestinal versus non-intestinal invasion events in L3 larvae and young adults. We found that L3 animals showed more invasion events in non-intestinal tissues compared to young adults. Around 59% of all sporoplasms were found in non-intestinal tissues when animals were at the L3 stage, while 31% of all sporoplasms were found in non-intestinal tissues when animals were young adults (Fig 5d). Similarly when we looked specifically at the muscle, on average 27% of sporoplasms in a given animal at the L3 stage were in the muscle, but this number decreases to 9% in young adults.

Together, these data support a model whereby sporoplasms may be directly delivered to both intestinal and non-intestinal tissue, for example by firing a polar tube from the lumen that can pierce all the way through the intestinal cells to the other side. Another possible model is that *N. displodere* sporoplasms initially invade intestinal cells, but then move independently of the polar tube to traverse the intestinal cell and reach other tissues as infection progresses. However, we were unable to find any evidence that sporoplasms move from one tissue to another. In fact, we found that sporoplasms were found in non-intestinal tissues as early as 2 minutes post-infection (S10 Fig), suggesting an incredibly rapid transit of the pathogen from intestinal lumen into non-intestinal tissue. In addition, we were unable to observe a significant increase in non-intestinal infection over time when L3 animals were pulsed with spores for exactly 1 hour and sampled immediately (1 hpi) versus 23 hours later (24 hpi) (S11 Fig), suggesting that if movement occurred from the intestine to non-intestinal tissues it would happen exclusively in the first hour of infection.

### *N. displodere* spores are not released continuously, but can exit when *C. elegans* burst

We next examined how newly differentiated *N. displodere* spores are released from infected animals. First, we quantified the production of *N. displodere* spores over the course of infection, measuring both the number of internal spores as well as the number of spores shed by infected animals. When we counted the number of spores inside intact (non-burst) *N. displodere*-infected animals from 4-10 dpi, we found a continuous increase in spore numbers over time such that by the last time point there was an average of 269,000 spores inside each animal (Fig 6a, *left*). By contrast, *N. parisii*-infected animals averaged many fewer internal spores, with a maximum of 34,000 internal spores per animal (Fig 6a, *right*). At these same timepoints, there were virtually no *N. displodere* spores shed into the media by these intact (non-burst) infected animals (Fig 6a, *left*), while *N. parisii* had a large number of spores being shed into the media at all timepoints in which internal spores were seen, with a peak observed at 6 dpi (Fig 6a, *right*). These results are consistent with our previous data showing that once new spores have differentiated, *N. parisii* has a continuous exit route from the *C. elegans* intestine by hijacking the host endocytic recycling pathway [21]. In contrast, *N. displodere* spores appear to have no continuous route of exit.

**Fig 6.**
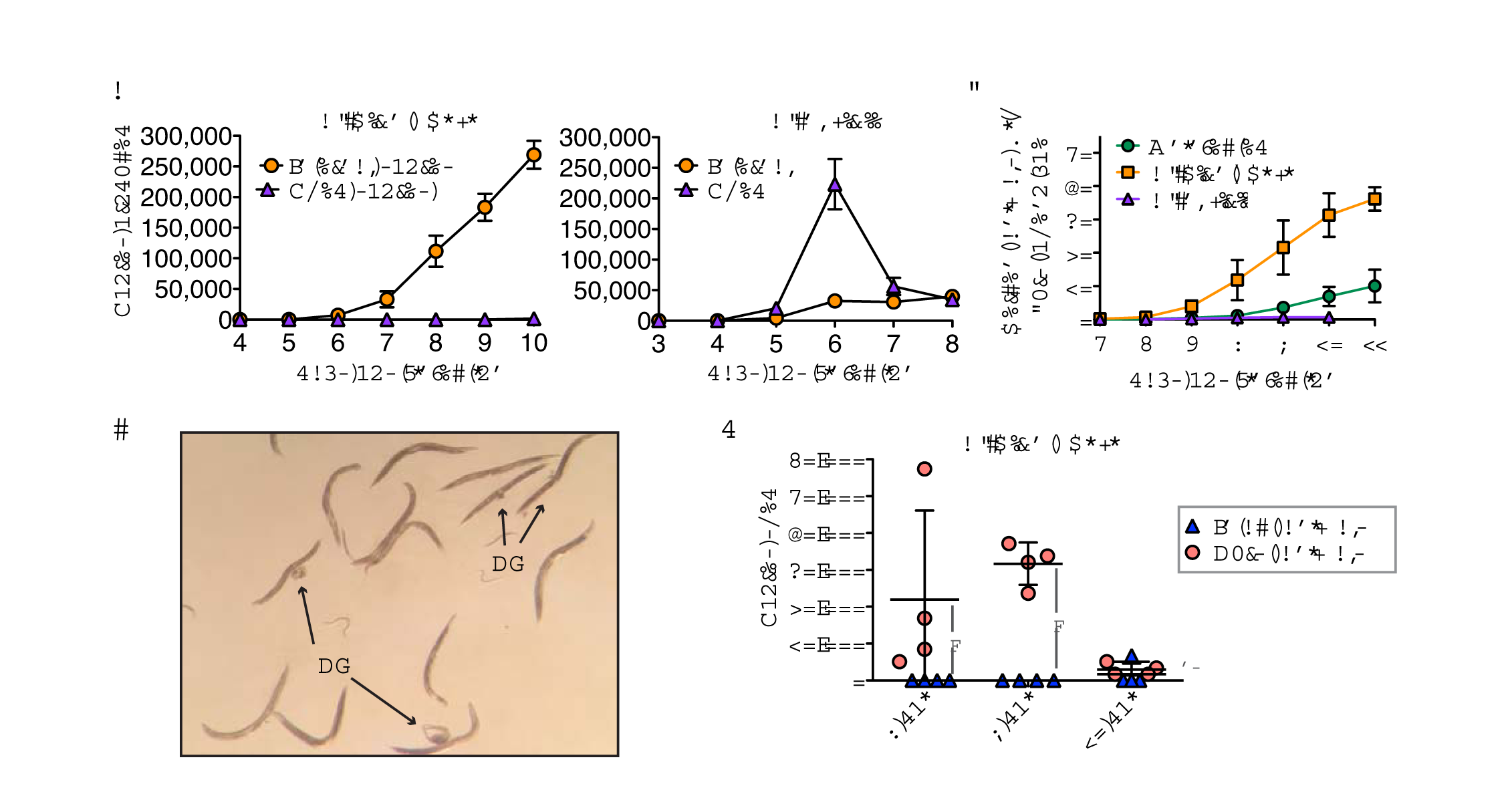
*N. displodere* spores exit through a bursting route. (a) Time course comparing the total number of internal spores compared to shed spores in *N. displodere*-infected (*left*) and *N. parisii*-infected (*right*) animals at 15°C. Note that only intact (non-burst) animals were picked for this assay. Internal spores indicate the average number of internal spores per animal, while external spores indicate the average number of spores shed by twenty animals into the media in four hours. Data points indicate the mean with SD of n=6 replicates of 20 animals across 3 experiments for internal *N. displodere* spores and n=4 replicates of 20 animals across 2 experiments for *N. displodere* shed spores and all *N. parisii* data. (b) Time course depicting the percent of animals with a burst vulva phenotype of uninfected, *N. displodere*-infected, and *N. parisii*-infected animals at 15°C. Data points depict the mean and SD from n=4 independent experiments for uninfected and *N. displodere* and n=3 experiments for *N. parisii* where each experiment consisted of triplicate samples containing at least 150 animals per replicate. (c) Image from a plate of wild-type *C. elegans* infected with *N. displodere* at 10 dpi. Indicated are adult animals with a burst vulva (*BV*) and internal organs spilling out. Image taken from a Nikon SMZ800 dissecting scope with an iPhone 5S. (d) Analysis of spores shed by late stage *N. displodere*-infected animals split into two groups, intact animals versus animals with a burst phenotype. Each data point indicates the number of spores shed by twenty animals for four hours of a single replicate, with the line and error bars showing the mean and SD of 4 replicates across 2 independent experiments (*p=0.0211, two-tailed Mann-Whitney test; ns=not significant, p=0.298).

To understand how *N. displodere* spores escape the host, we investigated a burst vulva phenotype seen in infected animals at late stages of infection. In this phenotype, the cuticle around the vulva breaks and internal tissues can be seen spilling from the opening, with the animal still alive and moving (S1 Video). By 11 dpi, 36% of *N. displodere*-infected wild-type animals displayed a burst vulva phenotype (Fig 6b). For reference, several infected animals with a burst vulva on a plate are shown (Fig 6c). By contrast, only 10% of uninfected and less than 1% of *N. parisii-infected* animals at this time point show this phenotype. Analysis of burst *N. displodere*-infected animals by microscopy shows that spores exit through this break in the vulva (see S1 Video, S12 Fig). In fact, when we tested for spore shedding in intact versus burst animals, we found that only burst animals shed *N. displodere* spores at 8 and 9 dpi (Fig 6d). By 10 dpi, the difference between these populations disappears, mostly due to a decrease in spores shed by burst animals. Additionally, these burst animals are infectious to new animals, while non-burst animals are not (S13 Fig). Based on this bursting phenotype, we have named this new microsporidian species *Nematocida displodere,* or ‘nematode-killer by causing to explode’.

### Analysis of the *N. displodere* genome

To investigate the genetic basis for traits displayed by *N. displodere* that distinguish this species from other *Nematocida* species, we sequenced, assembled, and annotated its genome. Assembly of the data resulted in a 3.066 Mb genome that is of comparable quality to other sequenced microsporidian genomes, both in terms of assembly statistics and the identification of proteins conserved throughout microsporidia (S1 Table). Phylogenomic analysis based on 87 single-copy orthologs present in 18 other sequenced microsporidia genomes and the outgroup *Rozella allomycis* revealed *N. displodere* to be a sister group to *N. parisii* and *N*. sp. 1 (Fig 7a). *N. displodere* proteins showed an average amino acid identity of 48.6% and 48.3% compared to *N. parisii* and *N*. sp. 1 proteins, respectively. For reference, there is 66% average amino acid identity between the proteins of *N. parisii* and *N*. sp. 1. The *N. displodere* genome is also smaller than the 4.148 Mb *N. parisii* (strain ERTm3) genome and the 4.700 Mb *N*. sp. 1 (strain ERTm2) genome [9]. This reduction is partly due to smaller intergenic regions in *N. displodere,* with 85.8% of this genome being protein coding, compared to 69.2% for *N. parisii* and 63.7% for *N*. sp. 1 (Fig 7b). Additionally, at 2278 predicted proteins, *N. displodere* has fewer proteins than either *N. parisii* or N. sp. 1, but shares 73.8% percent of its proteins with both species (Fig 7c,S1 Table, S2 Table). For comparison, there are 776 proteins that *N. par¡s¡¡*and *N*. sp. 1 share that are not found in *N. displodere,* but only 29 proteins that *N. displodere* shares with one *Nematocida* species that are not found in the other.

**Fig 7.**
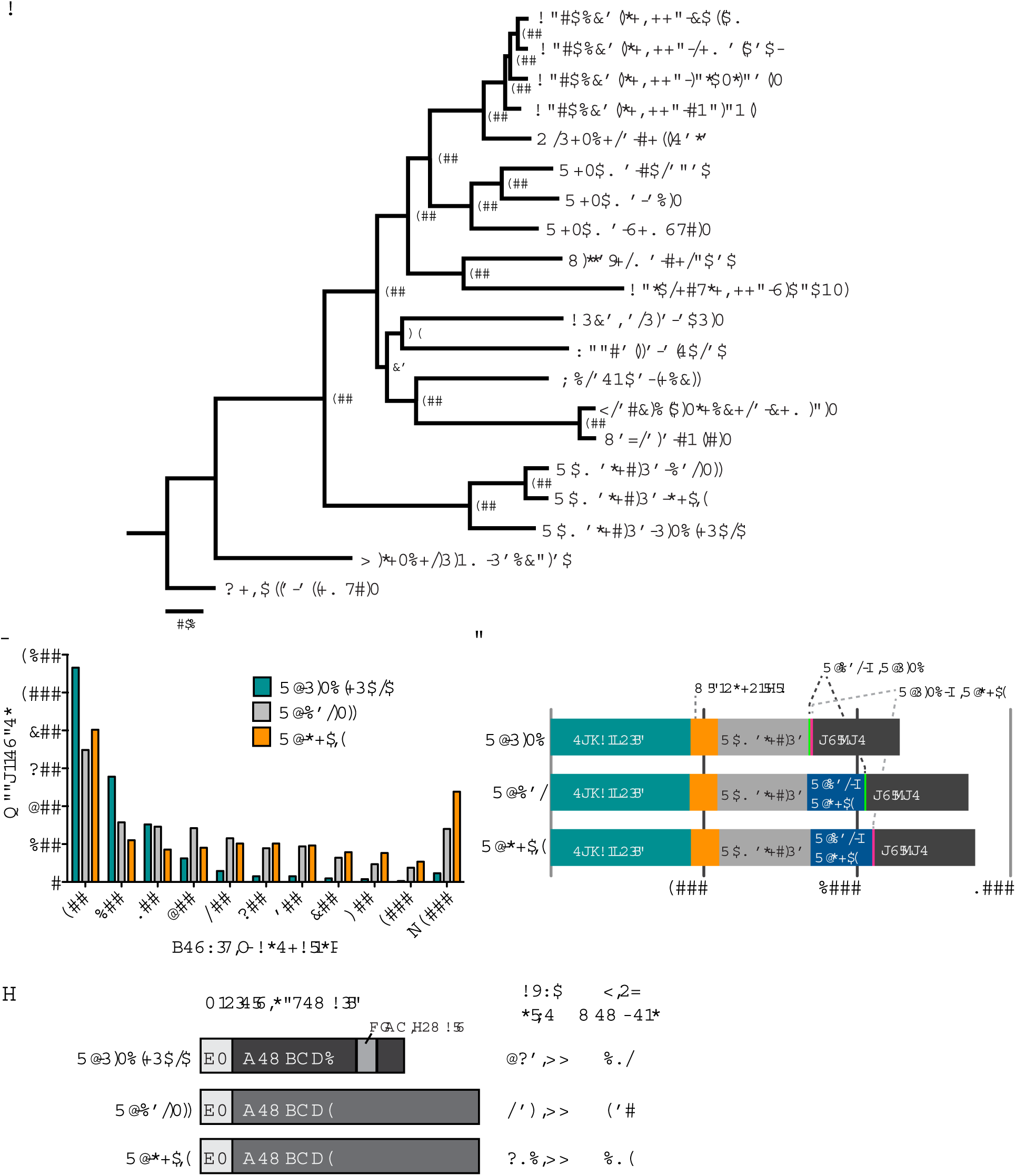
Analysis and comparison of *N. displodere, N. parisii, and N*. sp. 1 genomes. (a) Phylogenomic tree of *N. displodere* and 18 other microsporidia genomes, with *Rozella allomycis* as an outgroup. Bootstrap support is indicated next to each node. Scale bar indicates changes per site. The tree was created with FigTree 1.4.2 (http://tree.bio.ed.ac.uk/software/figtree/). (b) Histogram of intergenic region lengths of the three *Nematocida* species. (c) Comparison of protein content among the three *Nematocida* species. Proteins were classified into 7 categories: proteins shared with all *Nematocida* and at least 1 other non-microsporidian eukaryotic species (*eukaryotic*), proteins shared between all *Nematocida* and at least 1 other microsporidian species (*microsporidia*), proteins shared between the three *Nematocida* species (Nematocida), proteins shared by *N. displodere* and *N. parisii,* proteins shared by *N. displodere* and *N*. sp. 1, proteins shared by *N. parisii* and *N*. sp. 1, proteins not in any other species (*unique*). (d) Protein schematic of a generalized member of each of the large gene families in the *Nematocida* species, which contain signal peptides (*SP*). The average size of the gene family and the number of proteins in each species is indicated at the right.

One of the most striking features of the *N. displodere* genome is the presence of a large, expanded gene family containing 235 members, of which only two members were found in the *N. parisii* (NEPG_01491, NEPG_01930) and *N*. sp. 1 genomes (NERG_01194, NERG_02097) (Fig 7d), but no members were found in any other sequenced microsporidia species. This large gene family, named *Nematocida* large gene family 2 (NemLGF2) comprises over 10% of the predicted protein coding genes in *N. displodere*. The characteristics of NemLGF2 include an average length of 467 amino acids, the presence of a predicted signal peptide or N-terminal transmembrane domain on 152 members, and the presence of a RING domain on 113 members. The *N. parisii* and *N*. sp. 1 genomes contain another large expanded family (named NemiLGF1) [9] with 170 and 231 members, respectively (Fig 7d, S3 Table), but no members of this family were detected in *N. displodere*. We also identified several likely cases of horizontal gene transfer in *N. displodere* including a bacterial formamidopyrimidine-DNA glycosylase (NFDG_02224), which is a base excision enzyme evolved in DNA repair, and a FAD binding oxidase (NEDG_00514). Phylogenetic trees of homologous proteins support the idea that both of these enzymes are of bacterial origin (S14 Fig). Additionally, we identified homologs of an N-acyl phosphatidylethanolamine-specific phospholipase-D in the *N. parisii* (NEPG_01645) and *N*. sp 1 (NERG_00761) genomes, which was absent from *N. displodere*. A phylogenetic tree of these homologs supports this enzyme being of metazoan origin, most likely being acquired from a nematode (S14 Fig 14). Thus, several features, including large species-specific gene families and distinct horizontal gene transfer events, appear to distinguish the *N. displodere* genome from the *N. parisii* and *N*. sp. 1 genomes

For a **taxonomic summary** of this new species, *Nematocida displodere,* see S1 File.

## Discussion

Our study describes a new *Nematocida* species of microsporidia that infects *C. elegans,* and has characteristics distinct from the other species in the genus described to date. We found that *N. displodere* has a broad tissue tropism, with the capacity to infect and replicate in the muscle, neurons, epidermis, intestine, coelomocytes, and seam cells. Compared to *N. parisii,* which only infects and replicates in the intestine, *N. displodere* showed preferential tropism for the epidermis and muscle over the intestine. Likely due to this difference in tropism, we also found that *N. displodere* has an unusual mechanism for newly differentiated spore exit via bursting, while new spores of *N. parisii* continuously exit through defecation. When we analyzed the genomes of these related *Nematocida* species for differences, we found that *N. displodere* has a greatly expanded family of proteins with a RING domain of which only a couple members were found in the intestinal-trophic *Nematocida* species, and conversely a separate expanded family was found in these other *Nematocida* species that was completely absent in *N. displodere*.

To our knowledge, *N. displodere* has the broadest tissue tropism yet seen in *C. elegans,* and is the first pathogen with the capacity to infect the neurons, muscle, or coelomocytes of *C. elegans. N. displodere* infection is dependent on *C. elegans* feeding, which is consistent with other invertebrate-infecting microsporidia which display two major pathways for infection, either through ingestion of spores or transovum/transovarial passage [32]. Once in the intestinal lumen, the polar tube of *N. displodere* can infect intestinal cells, but is also long enough to theoretically reach other tissues, more so in younger animals than older animals. Because our data shows that *N. displodere* invasion of non-intestinal tissues can occur very rapidly, and microsporidia are not known to have standard movement apparatuses, like cilia or flagella [22, 33], it is likely that the polar tube is used to access other tissues. One model of infection is that *N. displodere* spores fire their polar tubes in the lumen to invade any tissue, intestinal or nonintestinal, to which the polar tube has access. This 'general access' model could explain why larger tissues like the muscle and epidermis showed a higher percent of pathogen than the neurons or less frequent cells, like seam cell and coelomocytes (see Fig 3d), assuming relatively similar proliferation rates in non-intestinal tissues. By contrast, we found that the polar tube lengths of similarly sized *N. parisii* spores were three-fold shorter than *N. displodere,* and these measurements correlated well with the number of polar tube coils seen in TEM crosssections, as *N. displodere* spores had up to five coils (see Fig 3g) while small *N. parisii* spores had only one coil in prior TEM images [18]. A shorter polar tube likely limits *N. parisii* infection to shorter distances from the intestinal lumen compared to *N. displodere* infection. However, given that the distance from the lumen to the basolateral side of the intestine is variable within an animal, with some distances being as short as 2 *μ*m (see Fig 2a, 5a), polar tube lengths are not the only limiting factor for infecting non-intestinal tissues from the intestinal lumen. It is possible that spores must convey enough force on the polar tube upon germination in order to pierce through multiple membranes to reach non-intestinal tissues [31], or that specific proteins are required on the polar tube for interaction with tissue-specific host factor on a cell for polar tube entry and invasion [34, 35].

Strikingly, although *N. displodere* can successfully invade the intestine, we found that the majority of intestinal infection fails to thrive when compared to the epidermis and the muscle. One possible reason for this tropism is that competition among other microbes in the intestine has put evolutionary pressure on *N. displodere* to efficiently proliferate in non-intestinal tissue. Multiple examples of potential competition among microsporidian species have been described, including the ecological observation that when the honey bee-infecting microsporidian *Nosema ceranae* is introduced into an area it can almost completely displace other naturally occurring microsporidia [36]. In addition, there is the laboratory observation that different species of microsporidia of the gypsy moth can exclude or suppress the growth of each other in a particular tissue [37]. In *C. elegans* in the wild, there may be an increased competition for resources in the intestine compared to other tissues, as several distinct pathogens can naturally infect and proliferate in the intestine [17, 18, 25].

A potential mechanistic reason for the tropism of *N. displodere* is that there is distinct induction of or sensitivity to tissue-specific host defense responses. For *C. elegans,* different tissue-specific transcriptional responses to intracellular pathogens have been identified for the epidermis upon *D. coniospora* infection and epidermal wounding, and for the intestine upon *N. parisii* and Orsay virus infection [27, 38]. Components of these transcriptional responses have been shown to play a role in tissue-specific host defense against their respective pathogens, including anti-microbial genes induced in the epidermis and Skp1-Cullin-F-box (SCF) ubiquitin ligase components induced in the intestine [26, 27]. For the epidermis we found that *N. displodere* failed to induce a reporter gene representative of the epidermal response, but induced reporter genes representative of the intestinal response to a higher degree than *N. parisii*. These observations could be representative of differential induction of tissue-specific defense responses upon infection Another possibility is that *N. displodere* has greater sensitivity to ubiquitin-mediated clearance in the intestine, resulting in the failure of the majority of infection in this tissue [27]. Future studies will investigate the mechanistic basis on the host side for the distinct tissue tropism of *N. displodere*.

To investigate the genetic basis on the pathogen side for the distinct features of *N. displodere,* we sequenced its genome and compared it with the other, intestinal-trophic *Nematocida* species. *N. displodere* is the earliest known diverging member of the *Nematocida* genus, but shares over 70% of its proteins with both *N. parisii* and *N*. sp. 1. The most striking difference between the genomes is the presence of species-specific expanded gene families, NemLGF2 and NemLGF1, found in *N. displodere* and the intestinal-trophic *Nematocida,* respectively, which are completely absent in any other sequenced genomes. An impressive 10% of the coding sequences in the *N. displodere* genome, or up to 33% of the proteins not shared with *N. parisii* and *N*. sp. 1, belongs to NemLGF2. These gene families likely evolved after the divergence from the last common ancestor and expanded as they adapted to their respective host environments. Considering the fact that microsporidia are obligate intracellular pathogens, it is likely that evolutionary pressure from the host played a role in the expansion, as has been shown for expanded gene families in other microsporidia species [39, 40]. In fact, the proteins in these expanded gene families are likely secreted into *C. elegans* cells at some point in infection, as both NemLGF1 and NemLGF2 have a high percentage of signal peptides among their members. Additionally, almost half of the NemLGF2 proteins from *N. displodere* had a C-2 terminal RING domain, typically found in ubiquitin E3 proteins and function to bind to E2 ligases[41]. The RING domains in NemLGF2 may serve as protein-protein interaction modules in the host cells, allowing these proteins to bind to host proteins to perform some yet unknown function. It is intriguing to speculate that the NemLGF2 proteins may play a role in interacting with the host ubiquitin system, as it has been shown to play a role in response to *N. parisii* infection [27], and RING domains are found in ubiquitin ligases. Despite having a broader tissue tropism than *N. parisii* and *N*. sp. 1, *N. displodere* has a smaller genome and fewer predicted proteins, which runs contrary to expectations that a broader range might require more genes to adapt to growth in varying niches and avoid different defense responses. However, a related observation was recently made in a study of two microsporidian species that infect mosquitos, as the microsporidia with a broad host range, *Vavraia culici,* had a smaller genome than the specialist microsporida, *Edhazardia aedis,* known to infect only one mosquito species [42]. Altogether, the discovery and characterization of *N. displodere* presents a unique model system in *C. elegans* to study the mechanistic and evolutionary basis of pathogen tissue tropism.

## Materials and Methods

### Nematode sampling and isolation

Wild nematodes were sampled from the woods near the Viosne stream in Santeuil, France on September 30, 2014 using methods previously described [17]. Wild *Caenorhabditis* animals that looked ‘sick’ were individually picked to nematode growth media (NGM) plates seeded with *E.coli* strain OP50-1, as described [43], and incubated at 20°C. P_0_ adults were allowed to selffertilize to produce F1 progeny and these P_0_ adults were analyzed by light microscopy for infection. *N. displodere* (designated isolate JUm2807, ZooBank ID, urn:lsid:zoobank.org:act:35CF055F-C311-4D9B-BFF0-B7B09FC441E4) was found in the head of the P_0_ of wild *C. elegans* strain (designated JU2807), isolated from the rotting stem of an *Asteraceae* plant (GPS coordinates: 49.12165, 1.95101) containing a proliferating population of approximately 500 *C. elegans* of various stages.

### *N. displodere* spore preparations

*C. elegans* strain JU2807 containing *N. displodere* isolate JUm2807 was cleared of all bacterial and fungal contamination by thoroughly washing a starved population of infected animals with sterile H_2_O and incubating 1 h in 15 ml H_2_O. Animals were incubated 2 h in S-basal (50 mM potassium phosphate, pH 6.0, 100 mM NaCl, 5 /μg/ml cholesterol) containing 100 /μ/ml gentamycin, 50 /μg/ml carbenicillin, 50 /μg/ml kanamycin, 20 /μg/ml tetracyclin, and 50 μ/ml streptomycin Sodium dodecyl sulfate (SDS) was added to a final concentration of 1% and incubated for 15 m. Finally, animals were washed with H_2_O and plated on NGM plates containing 50 /μg/ml carbenicillin, 25 /μg/ml kanamycin, 12.5 /μg/ml tetracyclin, and 37.5 μ/ml chloramphenicol seeded with concentrated OP50-1 bacteria, and incubated at 15°C for 5 d.

*N. displodere* spores were prepared as previously described for *N. parisii* [20]. Briefly, *N. displodere* JUm2807 was cultured by expanding large-scale cultures of antibiotic-treated *C. elegans* JU2807, followed by mechanical disruption of the nematodes, and then filtering to isolate spores away from animal debris. Similar methods were used to make *N. par/sii*spore preparations using the isolate ERTm1 infected in *C. elegans* N2.

### *C. elegans* strains and maintenance

All *C. elegans* strains were maintained as previously described [43]. The intestinal GFP strain ERT413 *jySi21[spp-5p::GFP; cb-unc-119(+)] II* was made in this study using Mos1-mediated single-copy insertion (MosSCI) [44]. Additional strains used in this study include:

- ERT54 *jyIs8[C17H1.6p::GFP; myo-2::mCherry] X*

- ERT71 *jyIs14[F26F2.1p::GFP; myo-2::mCherry]* [27]

- OH441 *otIs45[unc-119p::GFP] V*

- HC46 *ccIs4251[myo-3::GFP-NLS, myo-3::GFP-MITO] I; mIs11[myo-2::GFP] IV* [45]

- OH910 *otIs77[ttx-3p::kal-1, unc-122p::GFP] II*

- AU189 *frIs7[nlp-29p::GFP, col-12p::dsRed] IV* [26]

- DA465 *eat-2(ad465)*

- ERT125 *dyn-1(ky51)* [46]

- CB1368 *daf-2(e1368)*.

### Fluorescent in situ hybridization (FISH)

FISH was performed as described using FISH probes to the small subunit rRNA conjugated to CAL Fluor Red 610 (CF610) or 5-Carboxyfluorescein (FAM), with slight modification [27]. For single-species infections, a mixture of *Nematocida-specific* probes MicroA-CF610 (CTCTGTCCATCCTCGGCAA), MicroC-CF610 (CAGAATCAACCTGGTGCCTT), MicroD-CF610 (CGAAGGTTTCCTCGGATGTC), and MicroE-CF610 (GTACTGGAAATTCCGTGTTC) were used at 2.5 μg/ml each, with hybridization at 46°C and washes at 48°C. As indicated, the chitin-staining dye direct yellow 96 (DY96) was added at 10 μg/ml to the hybridization buffer to stain microsporidia spores in the animals [21]. For co-infection, a *N. displodere-specific* probe Microsp1A-FAM (CAGGTCACCCCACGTGCT) and a *N. parisii-specific* probe MicroF-CF610(AGACAAATCAGTCCACGAATT) were used at 5 μg/ml each, with hybridization at 52°C and washes at 54°C

### *C. elegans* infections with microsporidia

*C. elegans* strains were infected on NGM plates with purified *N. displodere* spores (isolate JUm2807) or *N. parisii* spores (isolate ERTm1) as described previously [9]. All infections in this study were conducted using a standardized dose of *N. displodere* or *N. parisii,* defined here as 3.5 × 10^4^ spores per cm^2^ (calculated from infecting a 6 cm NGM plate with 1.0 × 10^6^ spores). *N. displodere* was capable of being continually transmitted within *C. elegans* for multiple generations at 15°C, so all experiments were conducted at this temperature unless otherwise indicated.

For kinetics of infection, synchronized N2 L1 larvae were infected with a standard dose and sampled at 1-7 dpi for small subunit rRNA FISH in 24 h increments. Fifty animals per replicate were analyzed for sporoplasms, meronts, sporoblasts, and spores at each time point For microscopy with the tissue-specific GFP expression lines, strains ERT413, AU189, HC46, and OH441 were infected with the standard dose of *N. displodere* as synchronized L1 larvae. Animals were fixed at 3 dpi and stained by small subunit rRNA FISH, or at 5 dpi and stained by rRNA FISH plus DY96.

For analysis of the tissue distribution of *N. displodere* infection, synchronized L1 larvae of strains ERT413, AU189, HC46, and OH441 were infected at 15°C with half a standard dose of *N. displodere* spores in duplicate and fixed at 3 dpi for small subunit rRNA FISH. A total of 50 infected animals for each replicate were analyzed by confocal microscopy for meronts or meront clusters in GFP-positive tissues or GFP-negative tissues. Meront clusters were counted once if they were in distinct areas of the animal and/or separated from another cluster by at least 10μ/m. The percent of meront clusters in the GFP-positive tissue was calculated based on the total number of meront clusters calculated for each replicate.

For analyzing the percent of animals with tissue-localized symptoms at different time points, synchronized L1 larvae of strains ERT413, AU189, and HC46 were infected in duplicate at 15°C with 1/2 a standard dose of *N. displodere* spores in duplicate. Note that for 1 dpi animals, the L1 larvae were infected at the end of the L1 stage, by first growing for 22 h at 15°C before infecting with *N. displodere* so that the animals were in the L3 stage by the end of the experiment and express enough GFP for analysis. Animals were fixed at 24 hpi and 144 hpi for 1 dpi and 6 dpi, respectively, and stained by small subunit rRNA FISH for 1 dpi and FISH plus DY96 (10/μg/ml) for 6 dpi. The 3 dpi animals were calculated from the tissue distribution experiment (see above). Animals were analyzed for sporoplasms at 1 dpi, meronts at 3 dpi, and DY96-stained spores at 6 dpi and then the percent of animals with the symptom in the GFP-positive tissue were calculated for fifty animals per replicate.

### Transmission Electron Microscopy (TEM)

TEM was performed at the Electron Microscopy Facility, Department of Cellular and Molecular Medicine, UCSD. Synchronized N2 L1 larvae were infected with a standard dose of *N. displodere* at 15°C and harvested at 6 dpi and 8 dpi, and an uninfected sample was collected at 6 dpi. Animals were fixed with 2% of paraformaldehyde, 2.5% of glutaraldehyde in 150 mM sodium cacodylate buffer (SC), and washed with 150 mM SC buffer. Samples were post-fixed 3 h in 2% osmium tetroxide in 150 mM SC on ice, washed in 150 mM SC followed by ddH_2_O, pelleted in 2% agarose, and incubated in 2% uranyl acetate overnight at 4°C. Samples were dehydrated on ice with a graded series of ethanol from 50% to 100%, followed with 50% ethanol/50% acetone for 20 m, and twice in 100% acetone for 10 m. Samples were incubated in a graded series of Durcupan from 25% to 100% at RT. Finally, samples were incubated in 100% Durcupan ON at 60°C. Blocks were cut on Leica microtome with a diamond knife to 60 nm sections and collected on 300 mesh grids. Digital images were collected on a Tecnai TEM (Field Emission Inc.) at 80 kv by using Eagle 4K digital camera.

### Tissue-specific reporter infections

Synchronized ERT54, ERT71, and AU189 L1 larvae were grown on OP50-1 for 20°C for 24 h and 15°C for 24 h to the L3 larval stage. Animals were split and infected in triplicate with a standard dose of either *N. displodere* or *N. parisii* for 20 h at 15°C with approximately 800 animals per replicate. Animals were harvested and washed with M9 buffer and loaded onto a COPAS Biosort (Union Biometrica™) to measure GFP fluorescence and time-of-flight (TOF) of each animal. Data was analyzed using the R package COPASutils [47], with GFP expression of ERT54 and ERT71 normalized to TOF and AU189 normalized to *pcol-12::dsRed*expression.

### Infection of feeding mutants

For the *eat-2* mutant, synchronized N2 and *eat-2* L1 larvae were grown at 15°C to the gravid adult stage for 3 and 4 d, respectively, and then infected for 24 h with *N. displodere* at 1/10 the standard dose (3.5 × 10^3^ spores per cm^2^) to reduce the number of infection events per animal. For the temperature-sensitive feeding mutant *dyn-1(ts),* synchronized N2 and ERT125 L1 larvae were grown to the adult stage at 20°C for 3 d. Animals were then shifted for 2.5 h to either 30°C to stop ERT125 pharyngeal pumping (as monitored on a dissecting scope) or 20°C as a control, and infected with five times the standard dose (1.75 × 10^5^ spores per cm^2^) for 30 m at the respective temperatures. FISH was conducted as above and sporoplasms were counted and localized using a Zeiss LSM700 confocal microscope with a 40x objective.

To test dauer infection, synchronized *daf-2(e1368)* L1 larvae were grown for 2 d at 25°C to initiate dauer formation. As a control, *daf-2(e1368)* and N2 L1 larvae were grown for 2 d at 15°C to the L3 stage. All animals were infected with the standard dose of *N. displodere* at 25°C for 18 h. FISH was conducted as described above, except all animals were fixed in 100% acetone for 10 m to permeate the dauer cuticle.

### Stage-specific infecticn with *N. displodere*

Synchronized L1 larvae of strains ERT413 and HC46 were grown in duplicate for 50 h at 15°C to reach the L3 stage or 90 h at 15°C to reach the young adult stage. Synchronized L3 larvae or young adults were infected with a standard dose of *N. displodere* for 30 m at 15°C, then fixed for rRNA FISH. Twenty-five infected animals were analyzed by confocal microscopy and the number of sporoplasms in the GFP-positive tissue was compared to the total number of sporoplasms in the animal. To measure intestinal widths, a single z-plane image of an animal was taken with the lumen visible, and the distance was measured from the basal-lateral side of one cell, through the intestinal lumen to the basolateral side of the opposing intestinal cell.These values were halved to give an estimate of the distance from the intestinal lumen to the basal lateral side of the intestine.

### *C. elegans* bursting assay

Synchronized N2 L1 larvae were infected or mock-infected in triplicate with the standard dose of *N. displodere* and *N. parisii* at 100-200 animals per 6 cm plate, and grown at 15°C for 11 d. Infected adults were transferred to new plates at 5-8 dpi to remove the F_1_ generation. At 5-11 dpi all animals from each condition were analyzed by a dissecting microscope for a bursting phenotype and removed.

### Spore shedding and production assays

Synchronized N2 L1 larvae were infected in duplicate with the standard dose of *N. displodere* and *N. parisii* and grown at 15°C for 10 d. For quantifying spores produced in the animals, 20 animals were picked into 1 ml PBS + 0.1% Tween-20 (PBS-T) at 4-10 dpi for *N. displodere* and 3-8 dpi for *N. parisii,* and washed three times with 1 ml PBS-T. Animals were lysed and the number of spores produced per animal was counted as described [48]. For quantifying spores shed by the animals, 20 animals were picked into 500 μl of a 1:10 OP50-1 and M9 mixture at 5-10 dpi for *N. displodere* and 4-8 dpi for *N. parisii*. Animals were incubated for 4 h at RT with rotation, and secreted spores were separated from the animals and counted as previously described [21].

### Measurement of spore characteristics

For measuring spore dimensions, *N. displodere* spores were stained by 1:100 dilution of CFW (Sigma) and imaged with a 100x objective on a Zeiss AxioImager M1 upright microscope. Spores stained by CFW were measured using the light microscopy image with ImageJ (NIH) by length and width. One spore was removed as an outlier in the analyses because its length was greater than 1.5 times the interquartile range above the third quartile.

*N. displodere* polar tubes were stained by suspending spores in 0.5 ml PBS containing 0.5 mM H_2_0_2_, incubating on silane coated slide at room temperature in a humid chamber for 4 h, and adding 5 μL of 5 mg/mL NHS-succinyl-Rhodamine (Roche) for 1 h in the dark. *N. parisii* polar tube lengths were stained by subjecting *N. parisii* spores to two cycles of freeze-thaw, incubating on slides at room temperature for 4 h in PBS, and adding 20 ng/mL Concanavalin A conjugated to fluorescein isothiocyanate (FITC) and 10 ng/mL Caclofluor white for 30 m at room temperature [49]. Spores were visualized by washing twice with PBS and imaging at 100x objective on a Zeiss AxioImager M1. We measured the lengths of polar tubes that were still attached to spores with ImageJ (NIH) if the entire polar tube was in frame.

### Genome sequencing, assembly, and analysis

Spores were made as described above and further purified using a 50% Percoll (Sigma) gradient. DNA was extracted using a MasterPure Yeast DNA purification kit (Epicentre Biotechnologies). DNA was further purified using a DNeasy column (Qiagen). Genomic DNA sequencing data was generated using the MiSeq sequencing platform (Illumina), which resulted in 26,240,016 paired-end reads of 301 base pairs, resulting in ~2500X coverage. Sequencing reads were assembled into contigs and scaffolds using Abyss 1.5.2 with a Kmer value of 96 [50]. Only scaffolds and contigs of at least 500 bp were retained. Assembly statistics are in S1 Table. Gene prediction and orthology determination were done following procedures previously applied to other microsporidia genomes [9, 42]. This Whole Genome Shotgun project has been deposited at DDBJ/ENA/GenBank under accession LTDL00000000 for *N. displodere* JUm2807, and the version described in this paper is version LTDL01000000.

Genes were predicted from assembled scaffolds using Prodigal 2.60 [51]. Predicted proteins less than 100 amino acids were removed unless they had a PFAM match [52] of at least 10^-3^ or BLAST match against UniRef90 database [53] with an E-value of at least 10^-3^. A total of 2278 proteins were predicted. Each protein was assigned a standard name with the prefix NEDG. Proteins are listed in S2 Table.

Orthologous gene families were identified using OrthoMCL 2.0.9 [54] using an inflation index of 1.5 and a BLAST E-value cutoff of 10^-5^. Conservation of proteins for each microsporidian species was determined by counting the number of orthogroups conserved between all 19 species divided by the number of orthogroups conserved between the other 18 species. Phylogeny was constructed from 87 single copy orthologs present in *R. allomycis* and 19 microsporidian species (S1 Table). Proteins from each orthogroup were aligned using MUSCLE 3.8.31 [55]. These alignments were trimmed using trimAl 1.2 with the option-gappyout [56]. Each orthogroup alignment was then concatenated into a single alignment using FASconCAT 1.0 resulting in a total of 30,556 aligned amino acid sites [57]. ProtTest 3.4 was then used to determine that PROTGAMMALG was the best fitting model for the data [58]. Phylogeny was then inferred using the RAxML 8.2.4 with the PROTGAMMALG model and 1000 bootstrap replicates [59].

Categorization of protein conservation for the *Nematocida* species was done by identifying orthologous gene families with OrthoMCL. Six eukaryotes (*Saccharomyces cerevisiae, Monosiga brevicollis, Rozella allomycis, Neurospora crassa, Ustilago maydis,* and *Allomyces macrogynus*) and 19 microsporidia genomes were used (Table S1).

Large gene families were identified from OrthoMCL analysis. The proteins in these groups were used to build models of the families by aligning proteins with MUSCLE and building profile hidden Markov models using HMMER 3.0 with an 10^-5^ E-value cutoff [60]. The RING domain model for NemLGF2 was made by taking RING domains from NemLGF2 proteins and then searching for additional RING domains in NemLGF2 proteins. This process was iteratively repeated until no more domains with an E-value of at least 0.001 could be found.

Protein function was predicted with BlastKOALA [61]. PFAM domains in proteins were predicted with an E-value of 0.001. Signal peptides predicted with SignalP 4.1 [62], using the best model with a cutoff of 0.34 for both the noTM model and for the TM model. Transmembrane domains were predicted with TMHMM 2.0 [63].

Intergenic regions were calculated by subtracting the start of each coding gene from the closest preceding coding gene’s stop. Additionally the region before the start of the first predicted gene of a scaffold and the region after the stop of the last gene of the scaffold were included. Coding genes that were predicted to overlap were included and their intergenic value set to 0. Pairwise protein identities between species were calculated by aligning single copy orthologs with MUSCLE.

Putative cases of horizontal gene transfer were identified by BLAST hits against the NCBI non-redundant protein database, but not having a BLAST match with an E-value less then of 10^-5^ to any proteins encoded by the non-*Nematocida* microsporidia species listed in S1 Table. Both putative cases in *N. displodere* are in contigs that are bordered by genes that either have homology to a microsporidian protein or do not have detectable homology to any protein. The one putative case identified in *N. parisii* and *N*. sp. 1 is conserved between the two species and thus not likely to be contamination.

## Acknowledgements

A special thanks to Gaotian Zhang for helping in the early identification of the genus of *N. displodere*. We thank the *Caenorhabditis* Genetics Center, Jonathan Ewbank, Yishi Jin, Amy Pasquinelli, Sylvia Lee, and Andy Samuelson for *C. elegans* strains. Thanks to Lise Frezal for assistance with sampling and identifying wild nematodes. Thanks to Jimmy Becnel for his helpful comments on our TEM images. We thank Timothy Merloo and Ying Jones for preparing the TEM samples. Thanks to Robert Shoemaker for help with statistics and Reggie Harris for help with Latin and naming *N. displodere*. We thank Kirthi Reddy and Keir Balla for their helpful comments on the manuscript.

